# The Physics Network is Distinct from the Multiple Demand System in the Human Brain

**DOI:** 10.64898/2026.07.19.739417

**Authors:** RT Pramod, Sam Hutchinson, Nancy Kanwisher

**Author notes:** indicates equal contribution (RT Pramod, Sam Hutchinson).

## Abstract

Recent studies implicate regions of the human bilateral fronto-parietal cortices in intuitive physical reasoning—our ability to perceive, predict, and plan in the physical world. However, it remains unclear whether this "Physics Network" (PN) constitutes a distinct and domain-specific system, or whether instead it overlaps with the nearby domain-general multiple demand (MD) network. To answer this question, we scanned participants (N = 20) with fMRI in a pre-registered study and measured spatial overlap, response profile differences, and resting-state correlation between the PN and MD networks. We find that: (1) The PN and MD networks overlap only minimally within individual participants, (2) Each network exhibits distinct functional profiles across a broad array of tasks, and (3) Time courses of functional response are more correlated between subregions within the same network than subregions between networks. These findings indicate that the Physics Network is dissociable from the Multiple Demand system both spatially and functionally, supporting its distinct role in intuitive physical reasoning.

## Introduction

Our ability to perceive, predict, and plan actions in the physical world hinges on a sophisticated understanding of the physical structure of the world and the properties and behavior of objects within it. Recent neuroimaging studies have identified a set of fronto-parietal cortical regions—the ‘Physics Network’ (PN)—in the human brain that support this intuitive physical reasoning. The PN is activated preferentially for physical reasoning relative to difficulty-matched non-physical control tasks (Fischer et al., 2016). Recent studies have also shown that PN carries abstract information about object mass (Schwettmann et al., 2019), material properties (Paulun et al., 2025), the stability of configurations of objects (Pramod et al., 2022), and the presence of object-to-object contact (Pramod et al., 2025). Moreover, the PN represents predictions about imminent physical events, such as collision between two objects (Pramod et al., 2025). Additionally, the PN is sensitive to violations of physical but not social expectations (Liu et al., 2024), suggesting it implements domain-specific predictions. These findings collectively support the hypothesis that the PN contains a probabilistic generative model of the physical world like those used in video games, thus serving as "the brain’s physics engine" (Battaglia et al., 2013).

But is the PN truly a domain-specific system in the human brain, or does it merely represent a facet of a more general-purpose network? One reason to suspect the PN may have a more general function is that its anatomical distribution in the parietal and frontal cortices (see Figure 1) resembles that of the famously general-purpose "Multiple Demand" (MD) network, which is defined by its domain-general response to a wide variety of cognitive "demand" (Assem et al., 2020; Duncan, 2010; Duncan & Owen, 2000; Fedorenko et al., 2013). If the Physics Network overlaps with the Multiple Demand Network, that would challenge our current understanding of both systems. The MD system would be shown to depend more on the content of information processed than previously thought, and the PN would be shown to be less functionally specific than previously thought. If instead the overlap between the two systems is minimal, that would support current views of the domain specificity of the PN and domain generality of the MD network.

**Figure 1:**
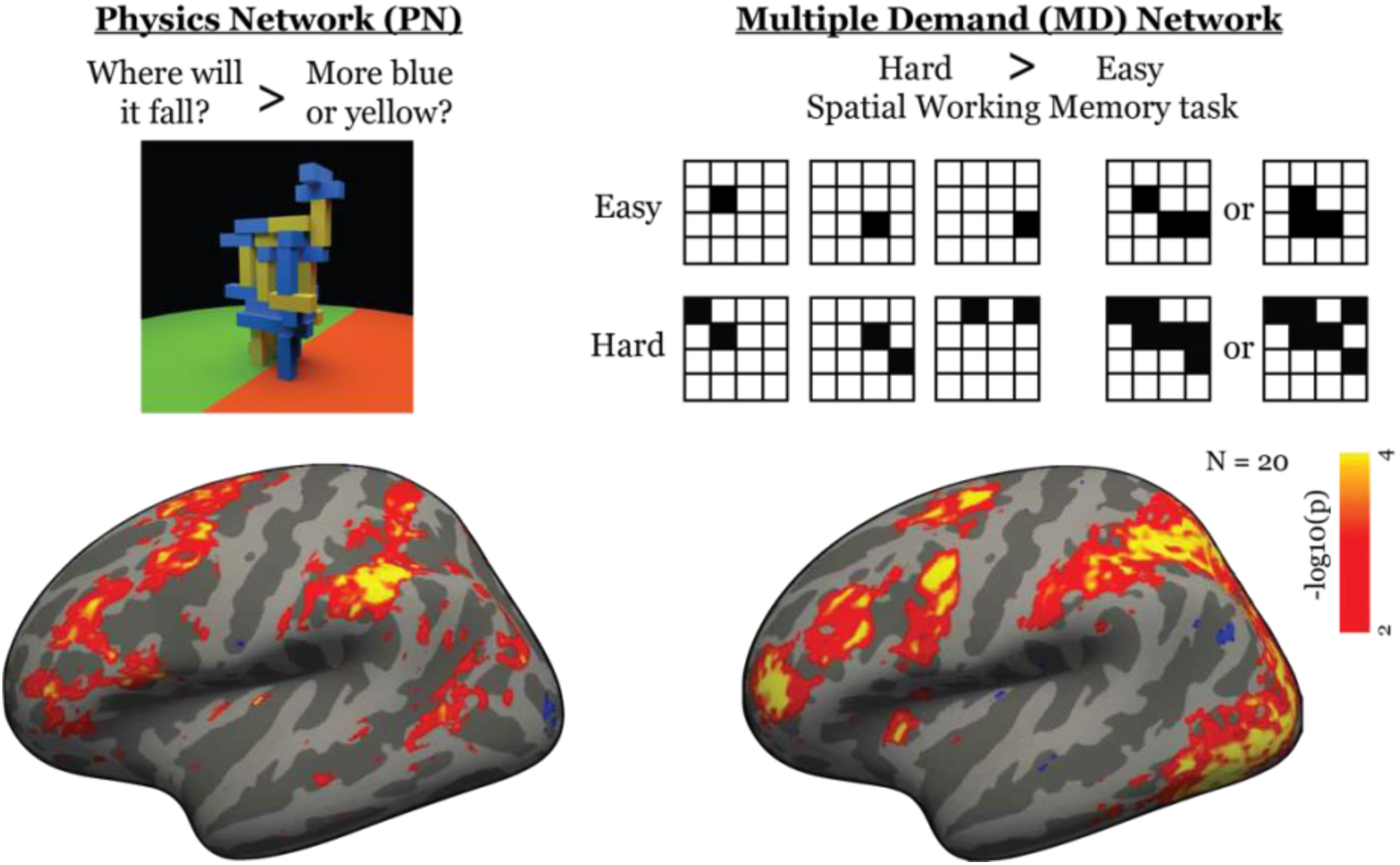
Group level random effects (rfx) analyses of the Physics Network (PN) and the Multiple Demand (MD) network. The rfx results are thresholded at p < 0.005 (uncorrected) and are displayed on the inflated surface of the left hemisphere of the human brain in the *fsaverage* space. The Physics Network was identified in each participant by the contrast between a physical and a color judgement task (top left). The Multiple Demand network was identified using the hard > easy contrast in a spatial working memory task (top right). Note that all the analyses presented in this study were performed in the volume space and surface maps are used for visualization only.

This question has been examined in a recent paper (Kean et al., 2025), which investigated spatial and functional overlap between the PN and the MD, language (Fedorenko et al., 2024), and theory-of-mind (Jacoby et al., 2016; Saxe & Kanwisher, 2003) systems in the human brain. That study revealed that the PN is spatially and functionally dissociable from language and theory-of-mind systems, but showed more complex relationships with MD. While the PN showed selective responses in the physics localizer contrast, some subregions of PN also responded strongly to the MD localizer contrast. A whole-brain Dice coefficient analysis revealed low-to-moderate overlap between PN and MD systems, suggesting only partial dissociation. However, this whole-brain analysis included regions outside the ‘core’ fronto-parietal PN (regions defined in Fischer et al., 2016), which may have biased the overlap estimates by including diffuse, weakly selective voxels.

To extend and refine these findings, we conducted a focused investigation of the core fronto-parietal PN and neighboring MD regions in individual participants. Our approach addresses three key limitations of prior work: 1) we focused specifically on the core fronto-parietal regions that have been most clearly implicated in intuitive physics, rather than measuring overlap across the whole brain, which includes regions activated by the standard physics localizer but whose role in intuitive physics is less clear; (2) we examined the function of both the PN and MD regions in detail, quantifying the response of each to 13 different stimulus and task conditions beyond the standard localizers; and (3) we computed resting-state functional correlations within and across networks to characterize their organization at the network level. This multi-level analysis— combining spatial segregation, functional selectivity, and network connectivity—provides a comprehensive test of whether PN and MD are truly dissociable systems.

First, we scanned 20 participants in order to localize each system. To identify the PN, we used the now-standard "Towers" localizer (Fischer et al., 2016; Figure 1), in which participants view videos of rotating tower of blocks, and judge either whether more blocks will land on the red or green side of the floor when the tower topples (the physics task), or whether the tower contains more blue or yellow blocks (the difficulty-matched non-physics control task). To identify the MD Network, we used a standard spatial working memory task (Fedorenko et al., 2013; Figure 1) in which participants view a sequence of two 4x4 grids, each containing either two (the difficult condition) or one (easy condition) filled-in cells, and then choose which of two sample grids contains all and only the filled-in cells from the sequence. The MD system is defined as the set of voxels that respond more strongly in the hard than easy version of this spatial working memory task. Critically, by running these two localizers in each participant we can ask whether the PN and MD networks overlap at the individual-participant level. This analysis is important to avoid potentially spurious overlap from group analyses that blur activations because of variability across participants in the precise anatomical location of each functionally defined region.

## Results

### Physics Network and Multiple Demand Network largely spatially segregate in individual brains

The apparent spatial overlap between the PN and MD group-level maps (see Figure 1) suggests a potential overlap between these two networks at the level of individual participants. However, anatomical variability of functional regions across participants can lead to spurious overlap in group analyses that is not reflected in individual participants. To test whether this is the case for the PN and MD network, we computed activation maps for the PN and MD localizer within each participant individually (see example individual-level maps in Figure 2A and for all participants in Figure S2) and then calculated the Dice-Sorensen coefficient for PN- and MD- selective regions in each participant. This metric quantifies the spatial overlap between two regions (ranging from 0 for no overlap to 1 for perfect overlap) while accounting for differences in their overall size (see Methods for details).

**Figure 2:**
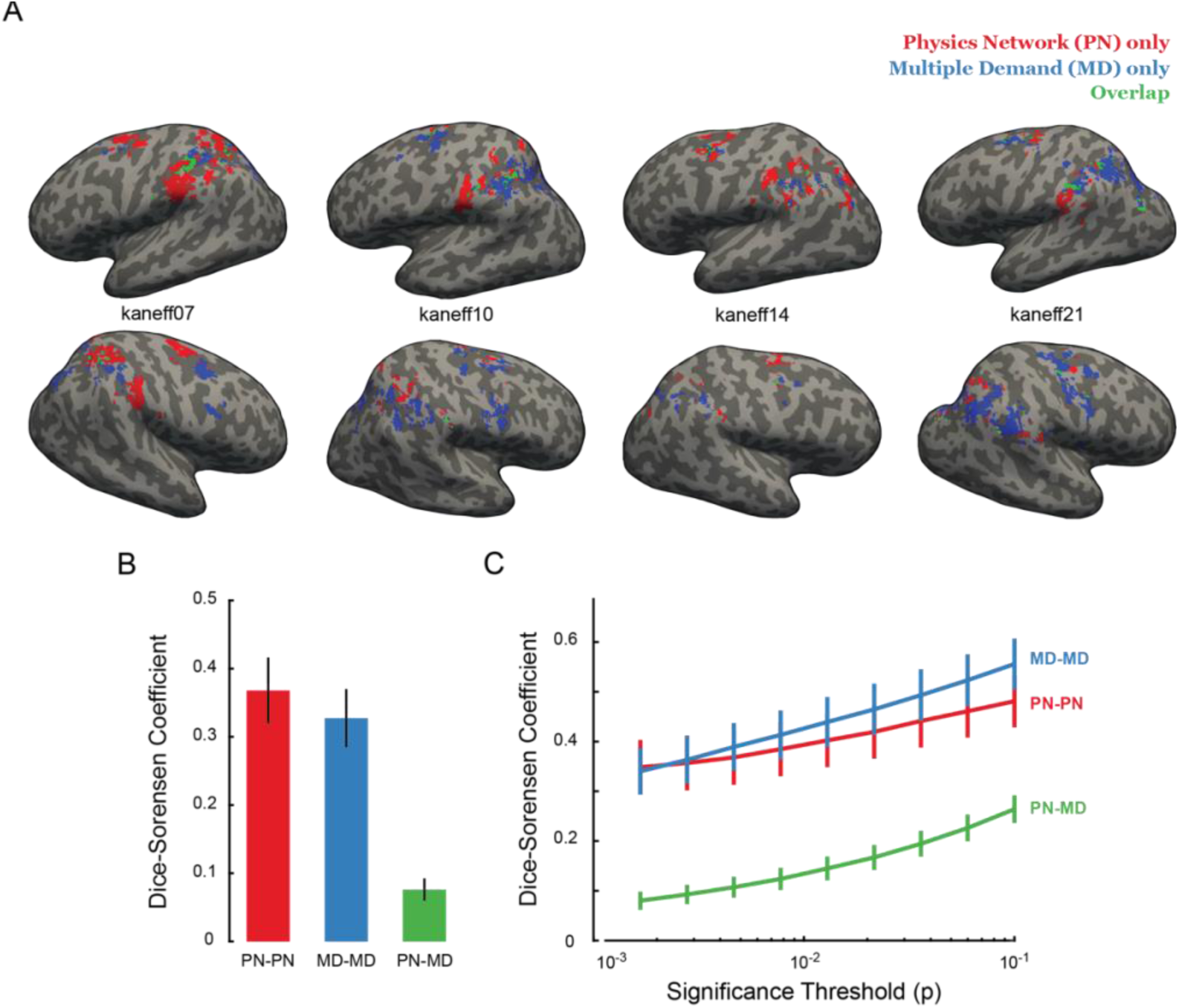
Dissociation between PN and MD networks in individual participants. A) Cortical surface projections showing PN- (in red) and MD-selective (in blue) vertices in the fronto- parietal cortices of 4 example participants. The vertices selective for both PN and MD are shown in green. The top and bottom rows show left and right hemispheres respectively. B) Dice- Sorensen coefficient computed over only significant voxels (p < 0.001) within-PN (i.e., across the two separate activation maps, each calculated based on half the PN localizer data), within- MD, and between PN and MD systems averaged across all participants. The error bars denote standard error of mean of the Dice-Sorensen coefficients across participants. C) Average Dice- Sorensen coefficient plotted as a function of significance threshold (p-value, in log scale) used to select voxels from each localizer contrast, showing the overlap between compared to within networks for all p-level thresholds. The error bars show standard error of mean across participants.

We found that the average Dice-Sorensen coefficient for overlap between PN and MD was 0.077 (standard error of the mean [s.e.m.] = 0.016 across participants), indicating very little overlap between these two networks at the individual level. In sharp contrast, within-network overlap— calculated by comparing independent splits of the localizer data—was substantially higher, with average coefficients (mean ± s.e.m. across participants) of 0.37 ± 0.05 for PN and 0.33 ± 0.043 for MD (Figure 2B). This difference was also significant as assessed in a repeated measure, mixed effect ANVOA with participants as a random factor and overlap type (within-PN, within-MD, and between-network) as a fixed factor (p < 0.00005 for the main effect of overlap type). Post-hoc analysis (using *multcompare* function in MATLAB) confirmed that both the within-network overlaps were significantly higher than the between-network overlap. Importantly, both within- and between-network overlaps were significantly greater than expected by chance (p < 0.0001; 10,000 bootstrap samples randomly assigning voxels within the combined parcel to either PN or MD, see Methods).

To examine whether spatial dissociation between the PN and MD networks differed across brain regions, we computed the Dice-Sorensen coefficient separately for parietal and frontal lobes. We found that the between-network overlap was lower than within-network overlap in both parietal and frontal lobes (only significant main effect of overlap type, p < 0.00005, in an ANOVA with overlap type and lobes as factors; Figure S3), indicating that the two networks are largely dissociable even within individual brain lobes. Thus, the spatial segregation between PN and MD is not limited to specific sub-regions but reflects a general organizational principle.

The spatial overlap results were also robust across different voxel selection criteria. The Dice-Sorensen coefficient for PN-MD overlap remained lower than within-network overlaps regardless of the statistical significance threshold (p-value) used to select voxels from the localizer contrasts (Figure 2C). Selecting the top 10% most selective voxels within each network produced similar results, with between-network overlap remaining low, and lower than within-network overlaps.

We further tested whether the spatial dissociation was specific to the spatial working memory task used to define MD. Using an alternative Math vs. Non-math contrast to localize MD (see Methods), we again observed stronger within-MD overlap (0.43 ± 0.034) than between-network overlap (0.073 ± 0.013; p < 0.00005 for the main effect of overlap type in a repeated measure, mixed-effect ANOVA with participants as a random factor and overlap type as a fixed factor; Figure S4), mirroring the results from the standard spatial working memory localizer. This consistency across different MD definitions strengthens the conclusion that spatial segregation between PN and MD is a robust feature of individual brain organization.

Together, these results establish that despite apparent overlap in group-level maps, the PN and MD networks are largely spatially segregated in individual brains.

### Physics Network and Multiple Demand Network show distinct selectivities

In the previous section, we provided evidence for the spatial dissociation between PN and MD regions. But two functionally defined regions can be spatially nonoverlapping even if they are functionally very similar, if one region is just over threshold for significance on one localizer contrast but just under significance for the other localizer contrast, and the other region shows the opposite pattern. Put another way, the spatial nonoverlap of two functionally defined regions is “a difference in significances, not a significant difference”. To statistically test whether the PN and MD regions in fact differ significantly from each other in function, and to more broadly characterize the functional selectivity of each, we first measured the response in each functionally defined region for the four conditions from the localizers in held-out data. Next, we did the same for 13 other stimulus and task conditions (see Methods). This analysis excludes the small set of overlapping voxels (which we examine separately below), allowing us to assess the functional profiles of each system in its core, selective regions.

First, when analyzing responses of each system to the four localizer conditions, we found a significant three-way interaction (p = 0.0001) in an ANOVA with functional regions (PN/MD), localizer task (Towers/Spatial Working Memory), and condition (test/control) as factors. Thus, the PN and MD regions indeed differed significantly from each other in their response to the four localizer conditions. Next, we analyzed each network individually.

In the PN (Figure 3A), as expected we found a significantly higher response to the Physics than Color condition in held-out data from the PN localizer (p < 0.0005 on a two-sided Wilcoxon signed rank test). We also found a significantly higher response to the Hard than Easy condition in the MD localizer (p < 0.0005 on a two-sided Wilcoxon signed rank test). However, the PN localizer contrast was significantly stronger than the MD localizer contrast (p = 0.039 for the interaction effect in an ANOVA on fMRI beta activations with localizer type [towers/spatial working memory] and condition [test/control] as factors). Thus, the PN-selective voxels respond to both task contrasts, but more strongly to the physics localizer contrast than to the spatial working memory task contrast.

**Figure 3:**
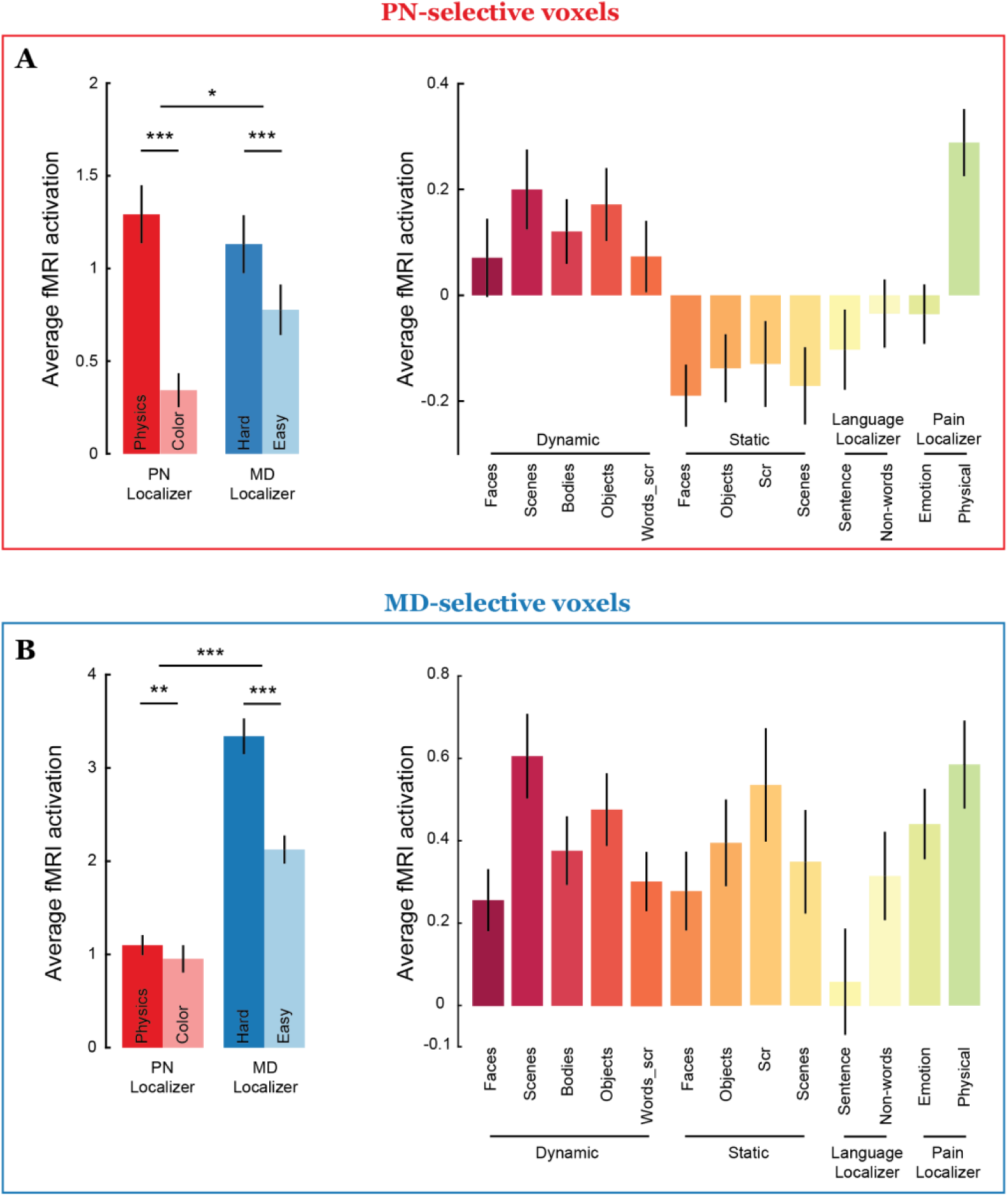
Functional dissociation between PN and MD networks. Average fMRI response magnitudes (betas) for all conditions in held-out data from the two localizer tasks and 13 other tasks in A) PN-selective voxels, and B) MD-selective voxels. The error bars denote standard error of mean of fMRI activations across participants. * is p < 0.05, ** is p < 0.005, and *** is p < 0.0005.

In the MD-selective region (Figure 3B), we again found significant effects of both the PN and MD localizer contrasts (Hard > Easy activations across participant: p < 0.0005; Physics > Color activations across participants: p = 0.0051 on a two-sided Wilcoxon signed rank test). However, the pattern was reversed: an ANOVA on fMRI beta activations revealed a significant interaction effect (p = 0.008), with the MD localizer contrast significantly stronger than the PN localizer contrast. This finding demonstrates functional selectivity in the opposite direction—MD- selective voxels show a stronger contrast for the spatial working memory task than for the physics localizer task.

To test whether this functional dissociation depends on the specific spatial working memory task we used as our MD localizer, we repeated the analysis with an alternative contrast to localize MD: Math vs. Non-math (Marvi, Hutchinson et al., 2025). In the Math condition, participants solved arithmetic problems presented auditorily. In the Non-math condition, participants heard false-belief and false-photograph stories (from a Theory of Mind localizer) and judged whether statements about the stories were true or false. The three-way interaction remained significant (p < 0.005, Figure S5), with PN-selective regions still showing a stronger effect for physics contrast and MD-selective regions showing a stronger effect for the math contrast.

The small set of voxels showing significant activation for both towers and spatial working memory localizers displayed a different pattern. These overlapping voxels showed no significant interaction between localizer and condition in an ANOVA (p = 0.79 for Towers/Spatial working memory overlap; p = 0.06 for Towers/Math overlap; Figure S6). This lack of selectivity suggests these voxels either contain intermixed populations of PN- and MD-selective neurons—below the resolution of fMRI to distinguish—or genuinely mixed-selectivity neurons that respond to both physics and cognitive demand.

Having demonstrated the functional dissociation between MD and PN systems, we then set out to characterize the functional response profile of each system in more detail, by leveraging data from a prior study (Marvi, Hutchinson, et al., 2025; see Methods for details) that collected BOLD responses across four functional localizer paradigms, comprising 13 different conditions (videos of faces, objects scenes, bodies, and words from the dynamic ventral pathway localizer; images of Faces, Objects, Scenes, and Scrambled Objects; written sentences and non-word stimuli; and written stories of characters experiencing either physical or emotional pain). We computed the response of each system to each of these 13 conditions (see Figure 3A-B) and found a significant interaction of region (MD/PN) by condition in an ANOVA (p < 0.0005). Further ANOVAs run separately on the data from each of the four localizers found a significant interaction effect between region and condition (p < 0.0005 for video-based localizer, p < 0.0005 for image-based localizer, p < 0.05 for the language localizer, and p < 0.005 for the pain matrix localizer). These analyses demonstrate clear functional differences in the response profile of the PN and MD systems, most strikingly a much stronger response to dynamic than static visual stimuli in the PN but not MD and a significantly higher response to stories describing physical than emotional pain in the PN but marginally so in the MD network (p < 0.0005 in the PN and p = 0.03 in the MD network on a two-sided Wilcoxon signed rank test comparing responses to physical and emotional pain conditions). The latter shows that the PN also responds to verbal stimuli while maintaining its selectivity for physical content (physical > emotional pain). We are delving further into this important finding of modality-general responses in the PN in ongoing work.

Taken together, these analyses show that the PN and MD systems are not only spatially dissociable, but also exhibit strikingly different response profiles.

### Physics Network and Multiple Demand Network form dissociable resting state networks

So far, our results show that PN and MD networks are both spatially and functionally separable in individual brains. But do they form distinct brain networks as well? Although PN has been referred to as ‘network’ in prior literature, direct evidence for strong functional correlation between sub-regions of PN is lacking (although see Navarro-Cebrián & Fischer, 2022 for correlated activity between Anterior Cingulate Cortex and the PN). We therefore used resting-state functional correlations (Figure 4A; see Methods for details) to address this question.

**Figure 4:**
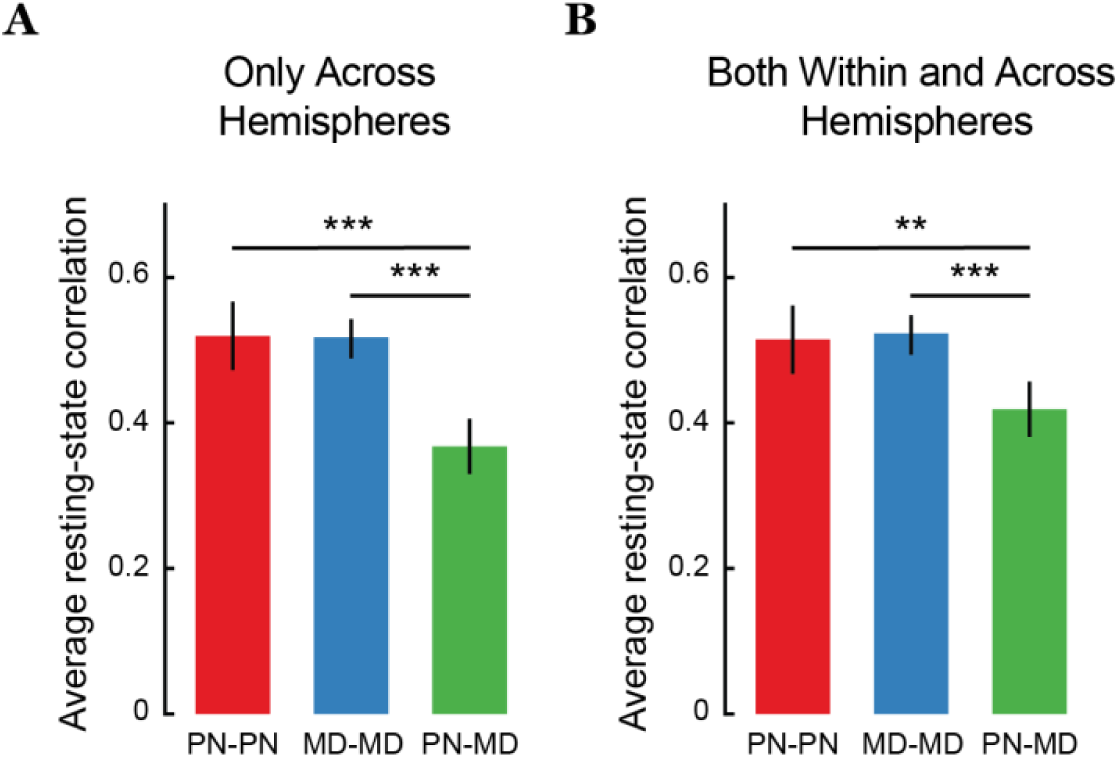
Network-level dissociation between PN and MD systems. Resting state correlations within PN (in red), within MD (in blue), and between PN and MD (in green) averaged over sub- regions and participants only across hemispheres (A), and both within and across hemispheres (B). The error bars denote standard error of mean across participants. * is p < 0.05. The statistical tests are performed on Fischer z-transformed correlations, but the plots show untransformed correlation values.

To unconfound cortical proximity from inferred connectivity, we computed within- and between-network functional correlations only across hemispheres (i.e., excluding ipsilateral correlations). We found that subregions within the PN network were correlated with each other during resting state (average functional correlation between subregions across left and right hemispheres: r = 0.52; Figure 4A). Similarly, subregions of the MD network in the fronto-parietal cortices were correlated with each other (average functional correlation between sub-regions across participants: r = 0.52). The correlations between subregions of PN were not statistically different from those within the MD network (p = 0.67 on a paired t-test), indicating that both networks exhibit similarly strong internal correlations. Critically, the functional correlation between subregions of different networks (i.e., PN and MD) was significantly weaker than correlations between subregions of the same network (average functional correlation between PN and MD subregions across hemispheres: r = 0.37; p < 0.0005 and p < 0.0005 for a paired t-test comparing against within-PN and within-MD correlations, respectively).

The trends in functional correlations remained consistent even when we included the ipsilateral comparisons: subregions within each network were strongly correlated with each other (average functional correlation: r = 0.52 and 0.53 for PN and MD respectively; Figure 4B) and this within-network correlation was significantly stronger than between-network correlations (average r = 0.42 between PN and MD sub-regions; p < 0.005 and p < 0.0005 for a paired t-test comparing against within-PN and within-MD correlations, respectively).

These findings provide the first evidence that PN constitutes a distinct cortical network with strong functional correlations between its subregions. More broadly, our results demonstrate that PN and MD form two separate cortical networks with distinct spatial and functional properties. Despite occupying similar anatomical territory, these networks maintain separate internal organization and distinct relationships with cognitive demands.

## Discussion

Despite observing substantial overlaps between group-level maps of the Physics Network (PN) and the Multiple Demand (MD) network in the human fronto-parietal cortices, we demonstrate in this study that these two networks are fundamentally distinct across three independent levels of analysis: i) they are largely spatially segregated at the individual level, ii) they show distinct functional selectivity profiles, and iii) they show greater resting functional correlation within each network than between networks.

At the individual-subject level, we found minimal spatial overlap between the PN and MD networks (Dice-Sorenson coefficient = 0.08), indicating substantial spatial dissociation despite the apparent overlap at the group level. This spatial segregation remained robust across different statistical thresholds and even within individual cortical lobes (see Supplementary figures S1-3). Using independent data to localize and measure network responses, we identified highly distinct functional selectivity profiles for the two networks (Figure 3) across a diverse set of 13 stimulus and task conditions. Most notably, this analysis showed a much higher response to dynamic than static visual stimuli in the PN but not MD network, and a highly selective response to stories describing physical versus emotional pain in the PN but less so in the MD network. This spatial and functional dissociation between PN and MD remained evident even when using alternative localizer tasks (e.g., Math task; Figures S4 and S5). Finally, resting-state fMRI revealed stronger functional correlations within each network than between them, indicating that the PN and MD participate in partly distinct networks (Figure 4). Our findings complement and extend prior work demonstrating partial spatial dissociation between PN and MD systems (Fischer et al., 2016; Kean et al., 2025), providing converging evidence from spatial, functional, and network correlation analyses that PN and MD are largely dissociable networks rather than overlapping subcomponents of a broader system. However, this functional segregation does not preclude interaction between networks: PN may communicate with the language network during tasks requiring linguistic description of physical events (the exportation hypothesis; (Casto et al., 2025; Liu et al., 2026)).

While our resting-state connectivity results demonstrate that PN and MD form dissociable functional networks, they do not address whether PN shares its cognitive load with other fronto- parietal networks. Our approach employed a functionally informed strategy: we first identified systems based on their responses to specific cognitive tasks, then examined their resting-state correlations. This approach has important advantages. By focusing on regions with specific functional selectivity, we can directly test whether systems that are functionally distinct are also distinct at the level of resting state networks. We also avoid the substantive interpretive challenge of reverse-engineering function from resting-state clustering. However, this functionally informed approach is necessarily constrained to networks with clear, measurable task responses. A complementary exploratory approach—applying resting-state clustering methods without prior functional information—could discover novel network organizations, reveal cross-network interactions that emerge only during rest, or identify networks specialized for functions not yet operationalized in laboratory tasks. We did not have enough resting state data to define these networks in individual participants reliably and hence we leave this comparison to future studies.

Thus, an important open question is how PN relates to the broader landscape of fronto- parietal networks. Prior work has characterized numerous resting-state functional networks in the fronto-parietal cortex like the Default Mode Network (Buckner & DiNicola, 2019; Du et al., 2024), the Dorsal Attention Network (Szczepanski et al., 2013), and the Fronto-parietal Network (Corbetta et al., 2008; Fox et al., 2005), involved in various cognitive processes including, attention, memory, planning, and mentalizing. Additionally, more functionally selective networks have been identified in nearby regions of fronto-parietal cortex, including the Action Observation Network (Caspers et al., 2010; Gazzola & Keysers, 2009; Goodale & Westwood, 2004), as well as networks supporting tool processing (Gallivan et al., 2013; Mruczek et al., 2013; Valyear et al., 2007) and numerical cognition (Harvey et al., 2013, 2015). An important open question is how PN relates to these other networks, particularly at the individual-subject level. Some theories propose that action and tool processing are computationally linked to physical reasoning, with PN forming an integral part of the brain’s embodied physics engine (Fischer & Mahon, 2021). Future studies employing complementary approaches—both fROI-based characterizations of action and tool usage and resting-state network approaches—could comprehensively map how physical reasoning integrates with other specialized and domain-general systems to support flexible goal-directed behavior in the physical world.

Our findings should be interpreted within the constraints of the neuroimaging and analyses methods used. Functional MRI has limited spatial resolution, preventing inferences about mixed selectivity at the level of single neurons (Rigotti et al., 2013). The small but significant spatial overlap observed between PN and MD could reflect two possibilities that fMRI cannot distinguish: either spatially intermixed populations of PN-selective and MD-selective neurons, or genuine populations of neurons that respond to both PN and MD task contrasts. Distinguishing between these alternatives would require higher-resolution imaging methods or complementary approaches such as electrophysiology.

Our results are robust to the specific task used to define the MD network. When we used a math task instead of the spatial working memory task to localize the MD network, we observed similar patterns of spatial segregation and even stronger functional dissociation between PN and MD (Figures S4 and S5). This consistency across task definitions suggests the dissociation is a robust feature of these networks rather than an artifact of our localizer choice. Nevertheless, future studies examining network dissociation using additional task manipulations could strengthen this conclusion. Experiments that directly contrast task difficulty with physical reasoning will be able to provide a more nuanced picture of spatial and functional dissociations between PN and MD within the fronto-parietal cortex.

Much remains to be learned about the neural basis of physical reasoning and its relationship to other neural systems. While we focused on the core fronto-parietal PN regions, additional regions beyond this core system also show selective responses to some physical reasoning tasks, including regions of the cerebellum (unpublished observations) and at least one region in the anterior frontal lobe (Kean et al., 2025). Another important question is how regions within the PN interact with other systems such as the language system (Casto et al., 2025) and systems for social cognition (Liu et al., 2026). Comprehensively mapping the brain systems supporting physical reasoning and understanding how these domain-specific systems interact with other neural systems remain an important challenge for future work.

## Methods

The methods and analysis plan were pre-registered and can be found here: https://osf.io/6vg2a/

### Participants

Participants were recruited from Massachusetts Institute of Technology and the surrounding Boston area (n=20, 10 females; aged 18-50). Before starting the experiment, all participants were given informed consent of the protocol approved by the Allen Institute for Brain Science and were subsequently compensated $75 for their involvement per two-hour scan session. The data presented in this paper were collected as part of a recent two-session study testing a new multi-function functional localizer (Marvi, Hutchinson et al., 2025).

### Stimuli and experimental design

#### Physics fROI Localizer (Towers task)

Each participant performed two runs of an ‘intuitive physics’ fMRI localizer task previously used to functionally define the frontoparietal physics engine in the brain (Fischer et al., 2016). In this task, subjects viewed short movies (∼6 s) depicting unstable towers made of blue, yellow and white blocks (see Figure 1) created using Blender 2.70 (Blender Foundation). The tower was centered on a floor that was colored green on one half and red on the other half such that it would topple toward one of the halves if gravity were to take effect. Throughout the movie, the tower remained stationary while the camera panned 360° to reveal different views of the tower. Subjects viewed these movies and were instructed to report whether more blocks would come to rest on the red or green half of the floor (‘physics’ task), or whether there are more blue or yellow blocks in the tower (‘color’ task).

Each run of this localizer task consisted of 23 18 s blocks: 3 fixation-only blocks, 10 blocks each of the physics and the color task. Each 6 s movie was preceded by a text instruction displayed on the screen for 1 s which read either ‘where will it fall?’ (‘physics’ task) or ‘more blue or yellow?’ (‘color’ task) and was followed by a 2-s response period with a blank screen. This sequence was repeated twice within a block with the same task cue but different movies. The subjects responded by pressing one of two buttons on a response box for each alternative in a task. The mapping of the buttons to the response was switched for the second run to rule out the effects of specific motor responses on the observed neural activations. We used a physics task > color task contrast to functionally identify the frontoparietal physics regions in each subject individually.

### MD fROI Localizer (Spatial Working Memory)

Each participant also completed two runs of a spatial working memory fMRI localizer task previously used to define the frontoparietal ‘multiple demand’ (MD) regions (Fedorenko et al., 2013). On each trial, subjects see a series of 4x4 grids and keep track of either four or eight locations (‘easy’ and ‘hard’ trials, respectively) which have been marked on these grids either one or two at a time (see Figure 1). At the end of each trial, subjects select which of two grids is showing the cumulative locations indicated on the previous grids in a two-alternative forced-choice decision.

Each run consisted of 48 eight-second trial blocks and four 16-second fixation blocks, with a palindromic ordering of hard and easy trials counterbalanced across runs. We used a hard > easy contrast to functionally identify MD regions in each subject individually.

### Efficient Localizer

In addition to the two localizers described above, each participant completed five runs of an efficient, multifunction localizer (EMFL; see Marvi, Hutchinson et al., 2025 for more details). The EMFL consists of five runs of ten 22-second stimulus blocks and three 18-second fixation blocks. Each stimulus block consists of a simultaneous presentation of stimuli from one visual category and one audio category. The visual stimuli are sets of short video clips showing dynamic examples from each of the following five categories: 1) faces, 2) scenes, 3) bodies, 4) objects, and 5) letters (superimposed over a scrambled-object backgrounds). The audio stimuli encompass the following five categories: 1) false belief stories and 2) false photo stories (presented in English), 3) arithmetic problems (also presented in English), 40 non-word sequences which were recognizably human speech, and 5) scrambled versions of the non-word sequences (‘quilted’ following the procedure of Overath et al., 2015). Each of these audio stimuli were broken into two segments: first, a *prologue* which sets up a scenario description (for the stories), lists a set of arithmetic operations (for the math), or establishes the gender of the speaker (for the non-word and quilted stimuli), and second, a *conclusion* which states a fact about the scenario (for the stories), gives an answer to the operations (for the math), or again makes the gender of the speaker obvious (for the non-word and quilted stimuli). Participants were instructed to answer a “matching” task by pressing one of two buttons in the scanner: a positive match occurs when the stated fact is logically consistent with the story (for the story conditions), the proposed answer to the arithmetic operations is true (for the math), or the gender of the speaker is the same for the two segments (for the non-word and quilted stimuli). There was no task related to the visual stimuli.

The stimuli were paired such that each of the 25 possible combinations occurs twice across the five runs, and each category appears twice and in a palindromic (for visual categories) or semi-palindromic (for the audio categories) order within each run. This allows us to disentangle responses from each modality separately and compute contrasts on the relevant conditions within each modality. We also collected data on a number of standard localizers designed to measure each of these contrasts separately (see Marvi et al., 2025 for details). This yields another 10 condition estimates from separate localizers. The condition profiles within PN and MD described in the results section above are vectors containing estimates of the responses to all 20 of the conditions described here (10 from the EMFL, 10 from the standard localizers).

To compare functional selectivity profiles between PN- and MD-selective regions, we chose only visual/written conditions (excluding auditory/speech conditions) across the localizers for a total of 13 conditions: 5 visual conditions from the efficient localizer, 4 visual conditions from the FOSS localizer (images of Faces, Objects, Scenes, Scrambled Objects), 2 conditions from the language localizer (written sentences and non-word stimuli), 2 conditions from the pain matrix localizer (written stories of characters experiencing physical or emotional pain).

### Data acquisition and preprocessing

All imaging was performed on a Siemens 3T MAGNETOM Tim Trio scanner with a 32- channel head coil at the Athinoula A. Martinos Imaging Center at MIT. For each subject, a high-resolution T1-weighted anatomical image (MPRAGE: TR = 2.53 s; TE = 3.57 ms; α = 9°; FOV = 256 mm; matrix = 256 × 256; slice thickness = 1 mm; 176 slices; acceleration factor = 3; 24 reference lines) was collected in addition to whole-brain functional data using a T2-weighted echo planar imaging pulse sequence. For 12 of the 20 subjects, the functional sequence had the following parameters: TR = 2 s; TE = 30 ms; α = 90°; FOV = 208 mm; matrix = 104 × 104; slice thickness = 2 mm; voxel size = 2 × 2 mm in-plane; slice gap = 0 mm; 52 slices. For the remaining eight subjects, to include better coverage of the cerebellum, we used a similar functional sequence with these parameters: TR = 2 s; TE = 30 ms; α = 90°; FOV = 208 mm; matrix = 104 × 104; slice thickness = 2 mm; voxel size = 2 × 2 mm in-plane; slice gap = 0 mm; *72 slices*.

Preprocessing was done using FreeSurfer (https://freesurfer.net/). All other analyses were performed in MATLAB 2015B (The Mathworks). fMRI data preprocessing included motion correction, slice time correction, linear fit to detrend the time series, and spatial smoothing with a Gaussian kernel (FWHM = 3 mm). Before smoothing the functional data, all functional runs were coregistered to the subject’s T1-weighted anatomical image. All analyses were performed in each subject’s native volume. The general linear model included the experimental conditions and 6 nuisance regressors based on the motion estimates (x, y, and z translation; roll, pitch, and yaw of rotation).

### Definition of individual fROIs

#### General Method

Functional regions of interest (fROIs) were defined using a group-constrained subject specific method (GSS; (Fedorenko et al., 2010; Julian et al., 2012)). Specifically, we used pre-defined parcels obtained from group-level probabilistic maps in prior publications to constrain the set of voxels for a given subject. We transformed parcels into the subject’s native space, and then selected voxels falling within these parcels based on their contrast significance in half of the localizer runs. Specifically, we selected voxels that showed significant contrast at p < 0.001 and if there were fewer than 50 voxels with this threshold, we reduced the threshold to p < 0.05. Subsequent analyses were performed in these selected voxels using functional data from remaining held-out runs. We also performed the spatial overlap analyses by choosing top-10% voxels within the parcel based on the contrast t-values.

### PN parcels

The GSS method was used to create the PN parcels in the physics localizer task. Specifically, we used the functional localizer contrast (physics > color task) data from 42 participants, all normalized to the fsaverage space, to create a probabilistic overlap map across individual subjects and across the whole brain. We then used watershed algorithm (‘spm_ss_watershed’) to obtain contiguous regions (or parcels) of voxels displaying physics > color contrast in at least 60% of the subjects. We then combined regions within the frontal and parietal lobes separately (based on Fischer et al., 2016) to create PN network parcels.

### MD parcels

Parcels for the MD fROIs (available at https://www.evlab.mit.edu/resources) were also constructed as a probabilistic overlap map of a large number of individual subjects’ (who also completed the spatial working memory localizer task) hard > easy contrast maps using the GSS method.

For all our analyses in this study, we combined PN and MD parcels within parietal or frontal lobes to obtain anatomical regions-of-interest (i.e., union of blue, red, and green voxels in Figure S1). *Note that all analyses in this study were performed in the volume space and the surface-based representations are only used for visualization*.

### Statistical analyses

#### Dice coefficient calculation

Once we defined our fROIs as described above, we computed the Dice-Sorensen coefficient as,

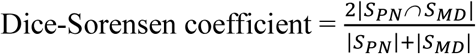

Where S_PN_ and S_MD_ are sets of voxels responding only to the physics localizer contrast and spatial working memory localizer contrast respectively, |.| operation denotes the cardinality (or, number of elements) in a set. We used the Dice-Sorensen coefficient to quantify the extent to which PN- and MD- selective regions overlap spatially in individual brains. Specifically, we computed the between-network overlaps using split-halves of the data: PN_even vs. MD_odd, and PN_odd vs. MD_even. We then averaged the two values to get an aggregate measure of the overlap. For within network overlaps, we computed the Dice-Sorensen coefficient for even and odd splits of the corresponding localizer task.

Note that we computed the Dice-Sorensen coefficient across the combined fronto-parietal PN and MD parcels. We show results for overlap within frontal and parietal parcels separately in supplementary materials (Figure S3).

### Bootstrap analysis

To ascertain how the observed overlaps compare to random chance, we computed bootstrap estimated overlaps from random sampling of voxels. Specifically, within the combined parcel, we uniformly randomly chose voxels and assigned them to either one network (or within-network split) or the other. We ensured that the number of voxels chosen this way matched the number of significant voxels observed in the original analysis. We then computed the Dice-Sorensen coefficient between the two sets of randomly chosen voxels. We repeated this analysis 10,000 times to get a null distribution of chance overlaps.

### fROI selectivity

We can further specify selective fROIs by defining the set of voxels that show a significant PN contrast but not a significant MD contrast (PN-selective), those which show the opposite pattern (MD-selective), and those which respond to both the PN and MD contrast. The crucial question of this investigation now becomes: do PN-selective voxels, identified using one split of the PN localizer (intersected with the combined PN and MD group level parcels), maintain their selectivity for physics>color and non-selectivity in the MD task (hard>easy) in independent data? Similarly, do MD-selective voxels (intersected with the combined PN and MD group level parcels) maintain their specificity for MD task and non-specificity for PN localizer in independent data? To assess this, we used the even (or odd) runs of each localizer experiment to select voxels that show a significant contrast response and then measured the responses of those same voxels in the odd (or even) runs of the experiment, within individual subjects. This allows us to obtain a set of eight measurements for each subject: the response to the physics, color, hard, and easy task conditions in PN- and MD-selective voxels. We can then submit these measures to a 3x2 ANOVA, where the factors are voxel set (PN- vs MD-selective), localizer (Physics localizer vs Spatial working memory localizer), and task condition (test condition [physics or hard] vs. control condition [color or easy]). We expected a significant triple-interaction effect between these factors indicating that voxels that only respond to the physics contrast but not the MD contrast, and vice-versa, in one split of the data cross-validate this selectivity in independent data.

To further investigate functional selectivity within PN- and MD-selective voxels, we compared activations in these sets of voxels in individual subjects to a set of independent task conditions (data from the efficient localizer in Marvi et al., 2025). These included a total of 20 conditions:

Dynamic Visual Faces, Scenes, Bodies, Objects and Words (static words against dynamic scrambled background); Static Visual Faces, Scenes, Objects and Scrambled Objects; Auditory False Belief, False Photo, Non-words, Quilted Speech and Math; Visual Sentences and Non-words from a language localizer; Auditory Non-words and Quilted speech from a speech localizer; and Auditory Physical Pain and Emotion Pain conditions. We split the data into even and odd runs in each individual subject and computed the average activation pattern for each split within PN- and MD-selective voxels. We then calculated Pearson correlation between even and odd splits of the data both within a network and also between networks.

A contrast of Math > (False Belief + False Photo) was used to define an alternate MD-selective network (Figures S2 and S3).

### Resting-state correlations

Having identified the PN- and MD-selective sets of voxels in each participant, we then calculated the correlation of their time courses during the 24 minutes of resting state fMRI (rs-fMRI) scanning. To do this, we first subdivided sets of voxels within each fROI based on their location in either the left or right hemispheres and frontal or parietal lobes. The parietal and frontal lobe parcels were composed of the union of PN and MD parcels within each lobe. We then averaged the time courses of all voxels which fall into each of the resulting eight sets (PN- and MD-selective x left/right hemispheres x parietal/frontal lobes), and then calculated the Pearson correlation for all pairs of these time courses. Finally, we average these correlations across subjects to yield an 8x8 similarity matrix. However, we retain only the upper or lower triangular matrix for further analyses as the 8x8 similarity matrix is symmetric. To assess the degree of within- versus between-network correlation, we computed the average correlation between subregions within one network and compared that to the average correlation between subregions of different networks.

