## Supplementary material for "The Physics Network is Distinct from the Multiple Demand System in the Human Brain"

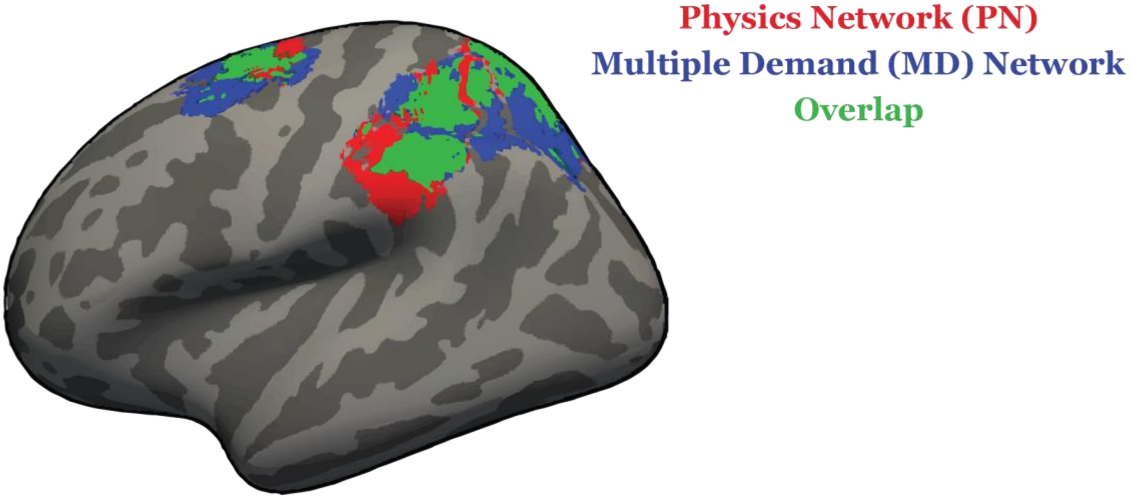


**Figure S1: Group-level parcels considered in this study for the Physics Network (red) and Multiple Demand Network (blue) shown on an inflated cortical surface.** The overlapping regions between the two networks is shown in *green*. The parcels were derived using the Group-constrained Subject Specific (GSS) method. For our functional region of interest (fROI) analyses, we used voxels from the union of PN and MD parcels shown here (i.e., all blue, red, and green voxels). Note that all analyses were done in the volume space, and surface maps are only for visualization.

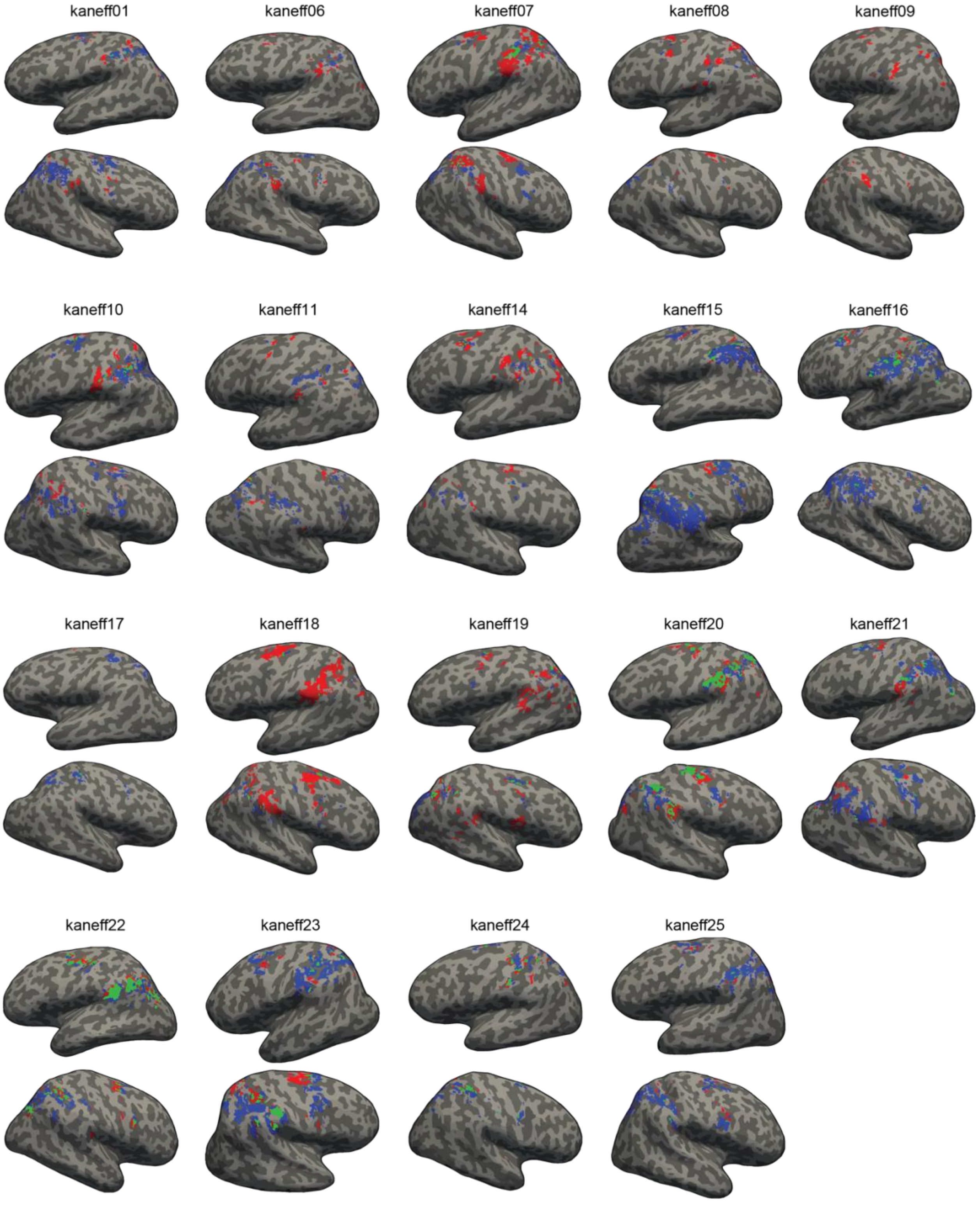


**Figure S2: Cortical surface projections showing PN- (in red) and MD-selective (in blue) vertices in the fronto-parietal cortices of all individual participants.** The vertices selective for both PN and MD are shown in green. Activation threshold was $p < 0.01$

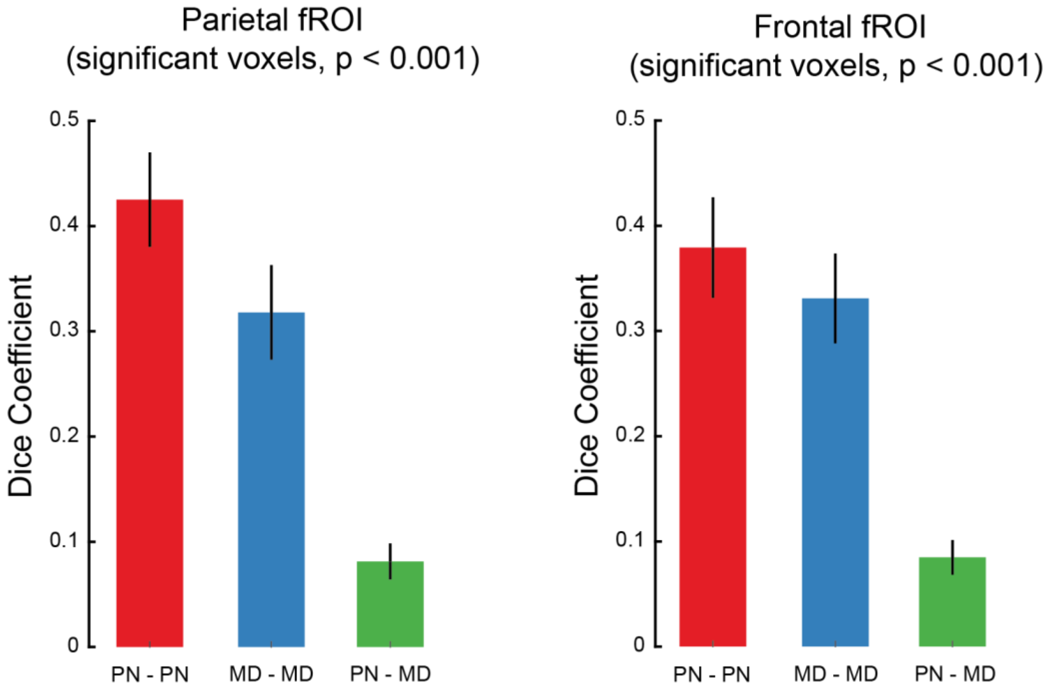


**Figure S3: Spatial dissociation between PN and MD in parietal and frontal cortices analysed separately.** The bar plots show Dice-Sorensen coefficient computed over only significant voxels ($p < 0.001$) in the Physics and Spatial Working Memory localizers. As in Figure 2B, the overlap coefficients were computed on two halves of the localizer data and averaged across all participants. In the parietal cortical fROI, we found that the average Dice-Sorensen coefficient for overlap between PN and MD was 0.083 (standard error of the mean [s.e.m.] = 0.017 across participants), indicating very little overlap between these two networks at the individual level. The within-network overlap—calculated by comparing independent splits of the localizer data—was substantially higher, with average coefficients (mean ± s.e.m. across participants) of 0.43 ± 0.045 for PN and 0.32 ± 0.045 for MD.

In the frontal fROI, the average Dice-Sorensen coefficient for overlap between PN and MD was 0.084 (standard error of the mean [s.e.m.] = 0.016 across participants). The within-network overlap was substantially higher, with average coefficients (mean ± s.e.m. across participants) of 0.38 ± 0.048 for PN and 0.33 ± 0.042 for MD.

An ANOVA on Dice-Sorensen coefficients with fROI (parietal/frontal) and overlap type (PN-PN/MD-MD/PN-MD) as factors revealed only a main effect of overlap-type ($p < 0.0005$) suggesting no significant difference between parietal and frontal fROIs in terms of network overlaps.

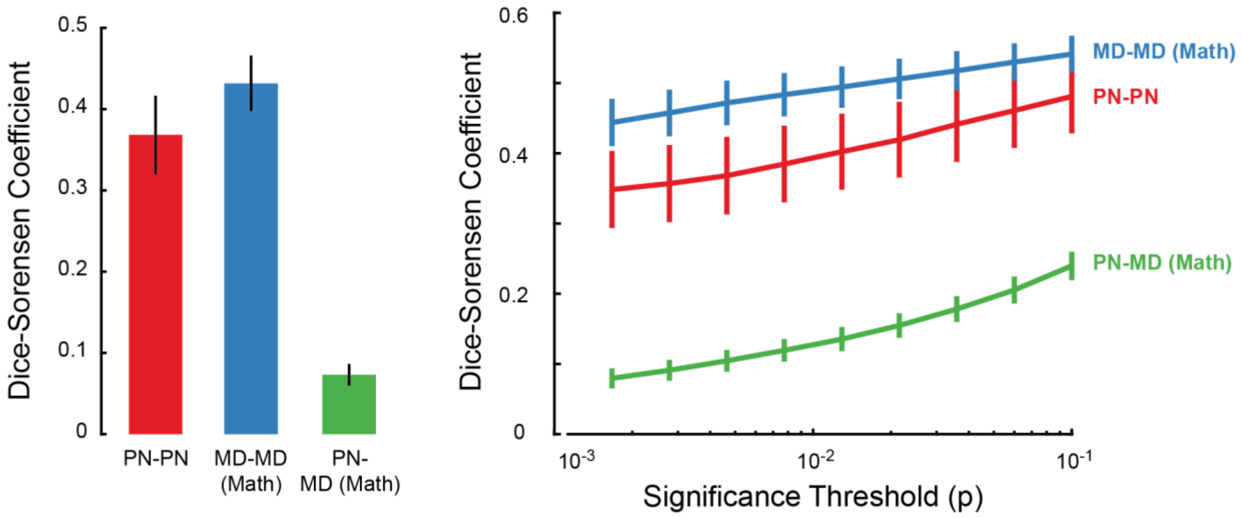


**Figure S4: Spatial dissociation between PN and MD using an alternate MD localizer.** The bar plot (left) shows the Dice-Sorensen coefficient computed over significant voxels ($p < 0.001$) within-PN, within-MD and between PN and MD systems averaged across all participants. The MD-selective voxels were obtained using the contrast Math > (False belief + False photo) in the efficient localizer paradigm described previously (Marvi, Hutchinson et al, 2025). The error bars denote standard error of mean of Dice-Sorensen coefficients across participants. We found that the average Dice-Sorensen coefficient for overlap between PN and MD was 0.073 (standard error of the mean [s.e.m.] = 0.013 across participants), indicating very little overlap between these two networks at the individual level. In contrast, within-network overlap—calculated by comparing independent splits of the localizer data—was substantially higher, with average coefficients (mean ± s.e.m. across participants) of 0.37 ± 0.05 for PN and 0.43 ± 0.034 for MD.

The line graph (right) shows average Dice-Sorensen coefficient as a function of significance threshold (p-value, in log scale) used to select voxels in each localizer contrast. The error bars denote standard error of mean across participants.

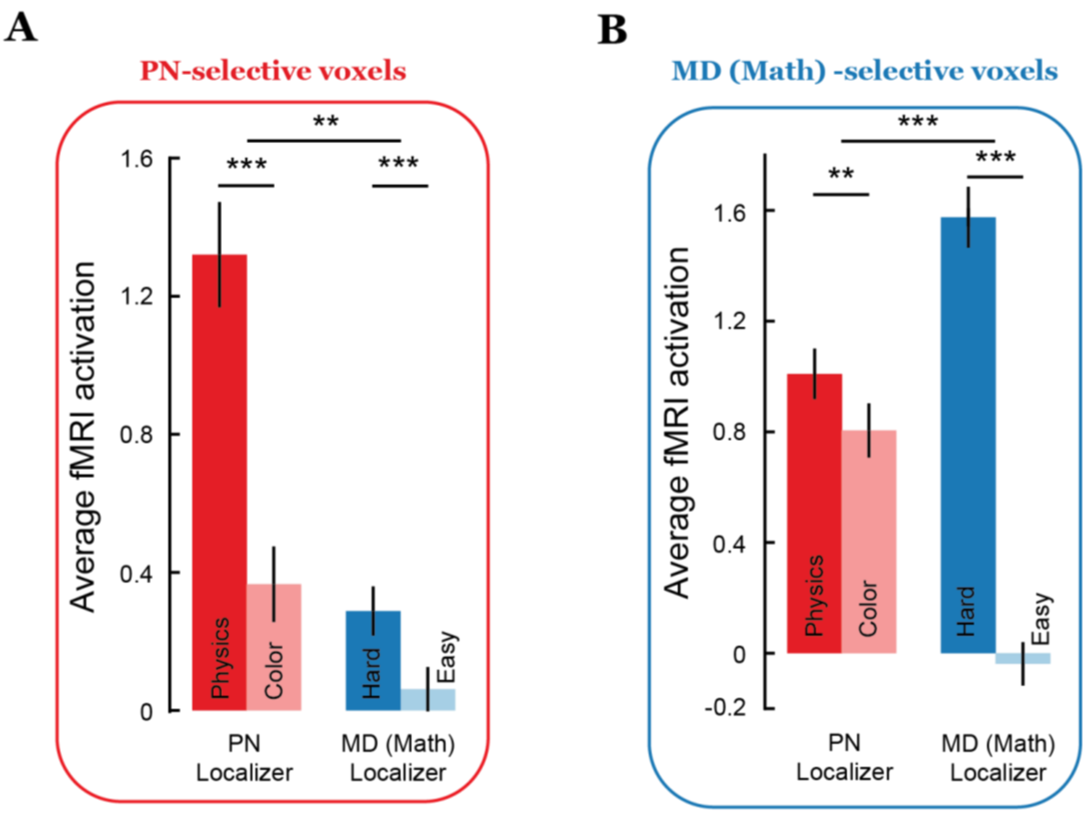


**Figure S5: Functional dissociation between PN and MD networks using an alternative MD localizer.** Average fMRI activations (betas) for all conditions in the two localizer tasks in A) PN-selective voxels, and B) MD-selective voxels using the Math > (False Belief + False Photo) contrast in the efficient localizer paradigm (Marvi, Hutchinson et al., 2025). The error bars denote standard error of mean of fMRI activations across participants. * is $p < 0.05$, ** is $p < 0.005$, and *** is $p < 0.0005$. In an ANOVA with functional regions (PN/MD), Task (Towers/Spatial Working Memory), and Condition (Test/Control) as factors, we found a significant 3-way interaction ($p < 0.0005$) indicating that selectivity profiles of the two regions are significantly different from each other.

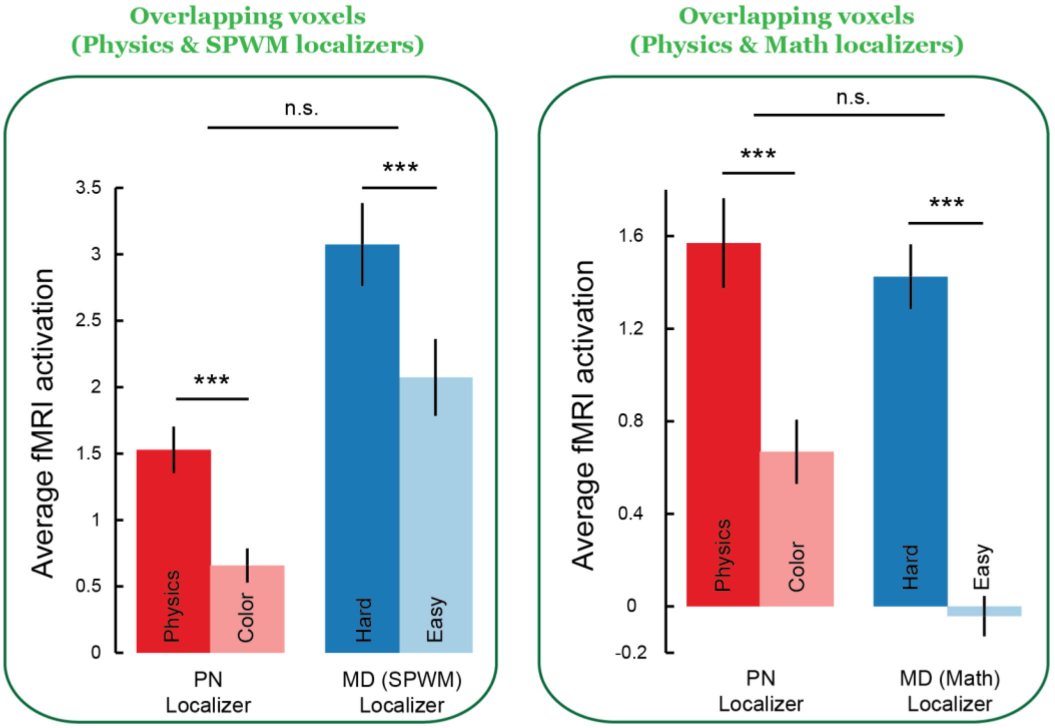


**Figure S6: Functional selectivity for overlapping voxels.**

For overlapping voxels across physics and spatial working memory localizers (left), we found in held-out data significant effects for both localizer contrasts (Average Physics > Color task contrast $p < 0.0005$ on a two-sided Wilcoxon signed rank test; Average Hard > Easy task contrast $p < 0.0005$ on a two-sided Wilcoxon signed rank test). An ANOVA on the beta activations across both localizers with localizer type (physics/spwm) and condition (test/control) as factors revealed no significant interaction ($p = 0.79$).

Similarly, for overlapping voxels across physics and math localizers (right), we found significant effects for both localizer contrasts (Average Physics > Color task contrast $p < 0.0005$ on a two-sided Wilcoxon signed rank test; Average Math > (False belief + False photo) task contrast $p < 0.0005$ on a two-sided Wilcoxon signed rank test). An ANOVA on the beta activations across both localizers with localizer type (physics/math) and condition (test/control) as factors revealed no significant interaction ($p = 0.06$).
